# Cultural and Ecological Dimensions of Spice Diversity: Indigenous Knowledge and Practices in Hadiya and Kambata-Tembaro, Central Ethiopia

**DOI:** 10.1101/2025.04.26.650802

**Authors:** Mulatu Osie

## Abstract

This study presents a comprehensive ethnobotanical investigation into the diversity and cultural significance of spices in the Hadiya and Kambata-Tembaro zones of Central Ethiopia. Through semi-structured interviews, focus group discussions, and participatory observations with local communities, including a strong representation of women were applied. Thirty five spice species, primarily herbaceous, with Lamiaceae emerging as the most dominant family documents. The findings underscore the centrality of homegardens as reservoirs of agrobiodiversity and hubs of traditional knowledge transmission. Spices were found to serve diverse roles culinary, medicinal, and ritual positioning them as biocultural keystone species intricately linked to local health systems, cultural identity, and daily life. Community preference rankings and culinary preparation methods revealed nuanced valuations based not only on taste but also cultural symbolism and perceived health benefits. Moreover, the study highlights gendered and socioeconomic dimensions of spice use and management, revealing persistent inequalities in resource access, decision-making, and benefit-sharing, despite women’s key roles in preserving and utilizing these resources. The research emphasizes the need for inclusive, gender-sensitive policies that recognize and protect indigenous ecological knowledge systems. It calls for participatory, community-led conservation strategies and further interdisciplinary studies on sustainable harvesting and the ecological dynamics of spice species across land-use systems. Overall, this study contributes to the growing body of knowledge advocating for the integration of traditional knowledge into national biodiversity and food security agendas, highlighting the importance of empowering local communities especially women to ensure the resilience of cultural and biological heritage amid environmental and socioeconomic change.

## 1. Introduction

Spices are integral to the culinary and medicinal traditions of Ethiopia, particularly in the southern regions where the Hadiya and Kambata-Tembaro peoples have long cultivated a rich diversity of plants. In these areas, spices such as garlic (*Allium sativum*), ginger (*Zingiber officinale*), and chili (*Capsicum frutescens*) are not only valued for their distinct flavors but also for their wide-ranging cultural and medicinal significance. These indigenous spices have been cultivated and utilized for generations, forming an essential part of the daily life and health practices of local communities (Wondimu et al., 2007). The intricate relationships between indigenous knowledge, agricultural practices, and the biodiversity of spice plants offer valuable insights into sustainable agricultural practices and the conservation of agro-biodiversity in the face of modern challenges such as climate change and agricultural expansion.

Homegardens, which combine domestic and agricultural activities in small, often irregular plots, serve as the primary landscape for spice cultivation in Hadiya and Kambata-Tembaro. These homegardens are known for their rich biodiversity, as they often host a variety of plants beyond just spices, including medicinal herbs and staple crops (Zemede Asfaw, 1997). The spatial structure and management of these gardens reflect adaptive strategies that help communities cope with environmental stressors and food insecurity, while simultaneously conserving valuable plant genetic resources (Eyzaguirre & Watson, 2002).

Indigenous knowledge of plant uses, particularly for medicinal purposes, plays a central role in the daily lives of Hadiya and Kambata-Tembaro communities. Spices such as *Aframomum corrorima* and *Ruta chalepensis* are commonly used in traditional medicine, addressing ailments ranging from digestive disorders to respiratory issues (Giday et al., 2009). This traditional ethnobotanical knowledge is passed down through generations, often orally, and is deeply rooted in the cultural fabric of these communities (Hunde et al., 2001). However, this knowledge is increasingly at risk of erosion due to factors such as urbanization, changing social structures, and the encroachment of modern agricultural practices (Adnan et al., 2020).

The diversity of spice plants in these regions also reflects the ecological complexity of the area, with microclimates and varying soil conditions contributing to a wide range of species growth. Studies have shown that regions with heterogeneous landscapes, such as those found in the Ethiopian highlands, tend to harbor higher plant diversity due to the presence of diverse ecological niches (Galluzzi et al., 2010). Understanding these ecological dynamics is essential for fostering sustainable practices that not only conserve biodiversity but also enhance food security and the resilience of local farming systems.

This study aims to explore the cultural and ecological dimensions of spice diversity in the Hadiya and Kambata- Tembaro Zones of Central Ethiopia. By examining the indigenous knowledge and practices surrounding spice cultivation, this research seeks to contribute to broader discussions on agrobiodiversity conservation and the role of traditional knowledge in sustainable agricultural systems. The study will also investigate the challenges and opportunities associated with preserving this valuable knowledge in the context of modern societal and environmental changes.

## 2. Materials and Methods

### 2.1. Study Area Description

This study was conducted in Hadiya and Kembata Tembaro Zones, two of the most densely populated zones in the Central Ethiopia Regional State. The study areas were chosen based on their rich history of spice cultivation and the prevalent use of homegardens for both agricultural and cultural purposes. The Hadiya Zone is characterized by an agroecological zone that includes highland and midland areas, while the Kambata-Tembaro Zone consists mainly of highland and plateau regions, providing a diverse array of growing conditions for spice cultivation.

Administratively, Hadiya Zone is subdivided into 17 districts, while Kembata Tembaro Zone comprises 14 districts. The administrative center of Hadiya Zone is Hosana, located approximately 267 km southwest of Addis Ababa, while Durame, the capital of Kembata Tembaro Zone, lies about 260 km southwest of the capital. According to population projections from the Central Statistical Agency (CSA, 2018), Hadiya Zone had a total population of 1,590,927, while Kembata Tembaro Zone had 902,073 inhabitants. The total land area of the zones is 3,593.31 km² for Hadiya and 1,355.90 km² for Kembata Tembaro.

#### Geographic Location and Environment

Both zones are situated along the fringes of the Great Rift Valley, giving rise to diverse topographical and agroecological conditions. Geographically, the study area lies between 7°05′ and 7°50′ N latitude and 37°15′ and 38°20′ E longitude, with elevations ranging from approximately 1,400 to 2,950 meters above sea level (Fig. 1). These altitudinal gradients contribute to variations in climate, soil fertility, and vegetation, fostering rich agro-biodiversity that supports a variety of crops, including spices and indigenous food plants.

**Figure 1.**
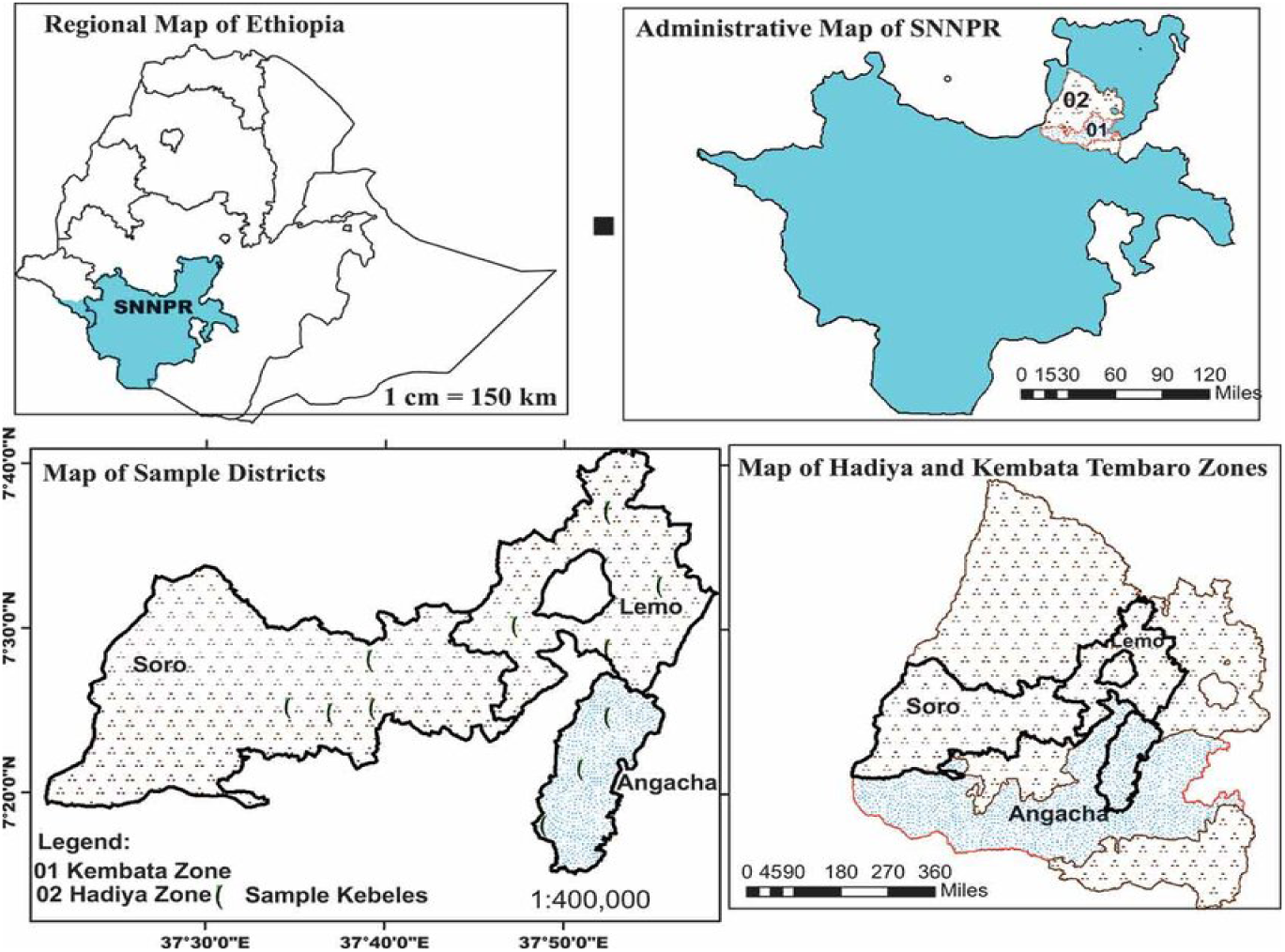
Map of the study area

#### Population and Demographic Characteristics

Together, Hadiya and Kembata Tembaro Zones are home to more than 2.4 million people, making them among the most densely populated areas in Ethiopia. The population is predominantly rural, with most households engaged in smallholder, rain-fed subsistence agriculture. Urbanization remains limited but is gradually expanding, particularly in key towns such as Hosana and Durame, driven by trade, education, and administrative services.

#### Cultural Landscape and Traditions

The cultural fabric of Hadiya and Kembata Tembaro Zones is a vibrant mosaic shaped by centuries of agrarian life, religious observance, and indigenous knowledge. Both communities are predominantly Protestant Christian, though Ethiopian Orthodox Christianity and Islam are also practiced. Annual celebrations such as *Meskel* (Finding of the True Cross), *Gena* (Christmas), and *Fasika* (Easter) are key religious and social events that reinforce communal bonds.

Traditional cuisine holds a central place in cultural expression. Dishes like *Kocho* (fermented enset bread), *Atmit* (spiced porridge), and various forms of *Wot* (stews) made with locally grown spices highlight the region’s culinary ingenuity. Food sharing during holidays and ceremonies not only reflects hospitality but also preserves the community’s ethnobotanical heritage and culinary identity. This dynamic interplay between traditional livelihood systems, ecological diversity, and cultural values makes the Hadiya and Kembata Tembaro Zones particularly important for studying indigenous plant use, spice diversity, and sustainable rural development within the broader context of contemporary Ethiopia.

### 2.2. Study Design

This study employed a robust mixed-methods framework to comprehensively investigate spice cultivation, conservation practices, and the embedded indigenous knowledge systems in the Hadiya and Kembata-Tembaro Zones of Central Ethiopia. Through the integration of ethnobotanical surveys, semi-structured interviews, and in-situ field observations, the research makes a substantive contribution to the preservation and documentation of traditional spice knowledge, a vital element of both cultural heritage and agrobiodiversity conservation. Employing a cross-sectional study design, the investigation captured the ecological and sociocultural dimensions of spice diversity across varied agroecological contexts. This approach facilitated the identification of spatial trends and variations in spice use, cultivation intensity, and local conservation strategies.

#### Sampling and Participant Selection

A purposive sampling strategy was employed to select eight kebeles (the smallest administrative units in Ethiopia) from each of the two study zones. Selection criteria emphasized kebeles known for their prominence in homegarden spice cultivation and their rich reservoirs of indigenous knowledge. Within each selected kebele, 48 households were randomly sampled, resulting in a total of 384 households (192 per zone). The sample predominantly comprised women, reflecting their central role and experiential knowledge in local spice cultivation, usage, and management. To ensure a comprehensive and gender-inclusive understanding of community practices, a smaller proportion of male participants was also included, allowing for the exploration of gender-specific roles, perspectives, and contributions to spice production, utilization, and conservation. This study was not retrospective in nature. It did not involve the use of archived samples, medical records, or previously collected survey data. All data were collected prospectively through direct interviews and observations during fieldwork focused on traditional spice use: (i) N/A– No retrospective data were accessed; (ii) N/A – No personally identifiable information was collected or accessed at any point during or after data collection. All participants’ identities were kept confidential.

### 2.3. Data Collection Methods

Data for this study were collected using a mixed-methods approach that integrated qualitative techniques to develop a comprehensive understanding of indigenous knowledge systems and community practices related to the use of spices and condiments. The primary data collection methods included semi-structured interviews, participatory observation, and focus group discussions (FGDs), each contributing unique insights into the cultural, culinary, and ecological dimensions of spice use. Additionally, preference ranking exercises were conducted to identify the most valued spices in traditional food preparation and to assess the major threats facing spices in the study area.

#### Semi-structured Interviews

Key informants were strategically selected to represent a broad spectrum of community knowledge and experience related to spice cultivation and use. These included experienced spice farmers with in-depth expertise in cultivation techniques, knowledgeable women recognized for their practical understanding of spice use in culinary and medicinal contexts, and elderly community members valued for their long-standing engagement with traditional practices and historical perspectives. Interviews were conducted using pre-designed guides that offered a flexible framework, enabling researchers to explore both predetermined themes and emergent insights.

The interviews sought to elicit detailed information on the diversity of spice species cultivated within the community; the multifaceted uses of spices, including culinary, medicinal, and cultural applications; traditional cultivation methods encompassing propagation, maintenance, and harvesting; and the indigenous knowledge systems guiding the conservation and sustainable management of spice resources.

All participants were adults and provided verbal informed consent, which was appropriate given the cultural context where written agreements are not customary. Consent was clearly explained, voluntary, and documented through detailed field notes. Community elders or local guides were often present as witnesses. No minors were included in the study, so parental or guardian consent was not required. As the study involved non-invasive ethnobotanical interviews, institutional ethics approval was not required.

#### Focus Group Discussions (FGDs)

Focus Group Discussions (FGDs) were conducted to gather collective knowledge and shared experiences related to spice cultivation and use. Each FGD consisted of 6 to 10 participants, purposefully selected to ensure representation from key social groups within the community. These included farmers, who offered practical insights into cultivation techniques and challenges; women, who shared their expertise in the preparation and domestic use of spices; and elders, who contributed historical perspectives and cultural knowledge.

The discussions focused on the community’s shared understanding of spice cultivation, the vital role of spices in sustaining local livelihoods, enhancing food security, and maintaining cultural identity. They also explored the environmental, economic, and social challenges affecting spice production, as well as the community-based strategies employed to conserve spice plant diversity and preserve associated traditional knowledge.

#### Preference Ranking

Preference ranking was employed to identify both the most culturally valued spices used in traditional food preparation and the most significant threats to spices in the study area. To this end, seven key informants were purposively selected to rank nine commonly used spices. A separate ranking exercise was conducted with the same informants to evaluate seven major threats to spice biodiversity. During this process, participants identified key challenges to spice cultivation, including land pressure, climate change, pest outbreaks, and market fluctuations. Additional data were also collected on local conservation practices, such as traditional preservation methods, seed saving, the use of organic fertilizers, and in-situ techniques like fencing and replanting.

Following the method outlined by Martin (1995), each informant ranked five to seven frequently encountered spices and threats by assigning integer values where higher scores indicated greater preference or perceived severity. The scores were then aggregated to produce an overall preference ranking for each spice species and threat factor. This approach enabled the identification of spices regarded as most important in cultural culinary practices and highlighted the most pressing threats to their sustainability within the local context. Furthermore, the Informant Consensus Factor (ICF) was calculated, based on the method proposed by Trotter et al. (1986), to assess the degree of agreement among participants regarding the medicinal applications of the spices.

#### Participatory Observation

Field visits were conducted to homegardens across each of the selected kebeles, providing opportunities for the direct observation of spice cultivation practices within their natural agroecological settings. These participatory observations enabled the systematic documentation of various aspects of homegarden management. Particular attention was given to the structural organization and spatial design of the gardens, the diversity of plant species cultivated with a specific focus on spices and the cultivation techniques employed by homegarden owners. Observations also included the conservation practices adopted to sustain spice diversity and productivity, such as organic inputs, mulching, and crop rotation. Furthermore, the spatial arrangement of spice plants was recorded, noting their integration with other crops and their role within the broader agroecological and household production systems.

#### Plant Identification

During field visits, plant specimens were collected from homegardens for taxonomic identification. Voucher specimens were prepared and initially identified using the *Flora of Ethiopia and Eritrea* (Edwards et al., 1997; Hedberg et al., 2009), and their identification was subsequently confirmed at the National Herbarium of Ethiopia, Addis Ababa University. The authenticated specimens were deposited in both the National Herbarium and the Botany Laboratory of Wachemo University.

#### Threats to Spice Diversity and Conservation Practices

To identify threats to spice biodiversity, participants were asked about challenges faced in spice cultivation, including factors such as land pressure, climate change, pests, and market demand. Data on conservation practices, including traditional methods of spice preservation, such as seed saving, organic fertilizers, and in-situ conservation techniques like fencing and planting, were also gathered through interviews and FGDs. The recruitment of participants took place between 15 February 2024 and 10 August 2024.

### 2.4. Data Analysis

Qualitative data obtained from interviews and focus group discussions (FGDs) were transcribed verbatim and analyzed thematically using NVivo software. Thematic analysis involved coding and categorizing responses to identify recurring patterns and themes related to spice cultivation practices, cultural significance, and indigenous knowledge systems. These themes were then interpreted to understand the social and cultural dimensions of spice use in the study communities.

Quantitative data were analyzed using SPSS software. Descriptive statistics were used to summarize socio- demographic characteristics and spice-related variables. Species diversity was assessed using ecological indices such as the Shannon-Wiener diversity index. Preference ranking data were analyzed to determine the most valued spices for culinary and medicinal uses. Informant Consensus Factor (ICF) values were calculated to evaluate the level of agreement among participants regarding the use of specific spices for various purposes, thereby identifying culturally important and widely accepted species. The integration of qualitative and quantitative analyses provided a comprehensive understanding of the ethnobotanical and sociocultural dynamics of spice use in the study area.

### 2.5. Ethical Statement

The study obtained verbal consent from adult participants, appropriate to the cultural context where written agreements are uncommon. Participation was voluntary, non-invasive, and documented through field notes with local witnesses. No minors were involved. Institutional ethical approval was not required due to the study’s focus on traditional spice use. Confidentiality was upheld, and the research adhered to recognized ethical principles, respecting indigenous knowledge and ensuring mutual trust.

## 3. RESULTS

### 3.1. Demographic Profiles of the Respondents

A total of 384 respondents (70 males and 314 females) were sampled for the study. The higher proportion of female respondents reflects the better knowledge and practice of female of spices in the study area.

Age distribution ranged from 20 to 67 years, with a mean age of 43.45 (±9.95 SD). In terms of education, 42.4% had completed primary school, while 33.6% had attained secondary education.

The average family size was 5.76 (±2.24 SD), with 89.1% of households having 4 to 7 members. The remaining 10.9% had 1 to 3 members. Although rare, the largest family size recorded was 9, observed in all study sites except Ashie.

### 3.2. Spices in the study area

A total of 35 spice and condiment species were documented in the Hadiya and Kambata-Tembaro zones, representing 27 genera and 17 botanical families. The species display diverse growth forms, with herbaceous plants comprising the majority (22 species, 63%), followed by shrubs (8 species, 23%) and trees (5 species, 14%). This dominance of herbaceous species reflects their accessibility, rapid growth cycles, and suitability for small-scale cultivation, which make them particularly prominent in both culinary and medicinal applications.

The most commonly used and culturally significant spices were predominantly herbaceous, reinforcing their centrality in the community’s ethnobotanical practices. Lamiaceae emerged as the most species-rich family with 8 representatives, followed by Apiaceae (5 species), Zingiberaceae (4), Rutaceae (3), and Asteraceae (2). The remaining families were represented by single species, collectively contributing to the region’s rich phytocultural landscape.

In terms of habitat distribution, the majority of species (28 species, 80%) were sourced from homegardens, emphasizing the role of household agroecosystems in conserving spice diversity. The remaining 7 species (20%) were obtained from cultivated fields, suggesting a supplementary role of formal agriculture in spice production. This pattern aligns with ethnobotanical trends observed in other Ethiopian regions, where homegardens serve as dynamic reservoirs of both biodiversity and traditional knowledge.

Notably, the local communities exhibit profound indigenous knowledge related to these spices, including detailed vernacular nomenclature, morphological identification, and multi-purpose applications spanning culinary, medicinal, and ritual domains. This highlights the importance of preserving both the biological and cultural dimensions of spice diversity as integral components of local heritage and sustainable livelihoods.

### 3.3. Diversity of spices

Our study revealed distinct patterns of spice species diversity across the investigated sites. Homegardens exhibited the highest Shannon diversity (2.62), indicating a richer and more diverse spice composition compared to farmland (1.68) and the wild area (0.64) (Table 3). Consistent with this, the Simpson diversity index also showed the highest values in homegardens (0.88), followed by farmland (0.69) and the wild area (0.42). The elevated Shannon diversity in homegardens appears to be linked to a more equitable distribution (higher evenness) of spice and condiment species within these systems, contrasting with the less even distribution observed in farmland and wild areas (Table 3).

**Table 1:**
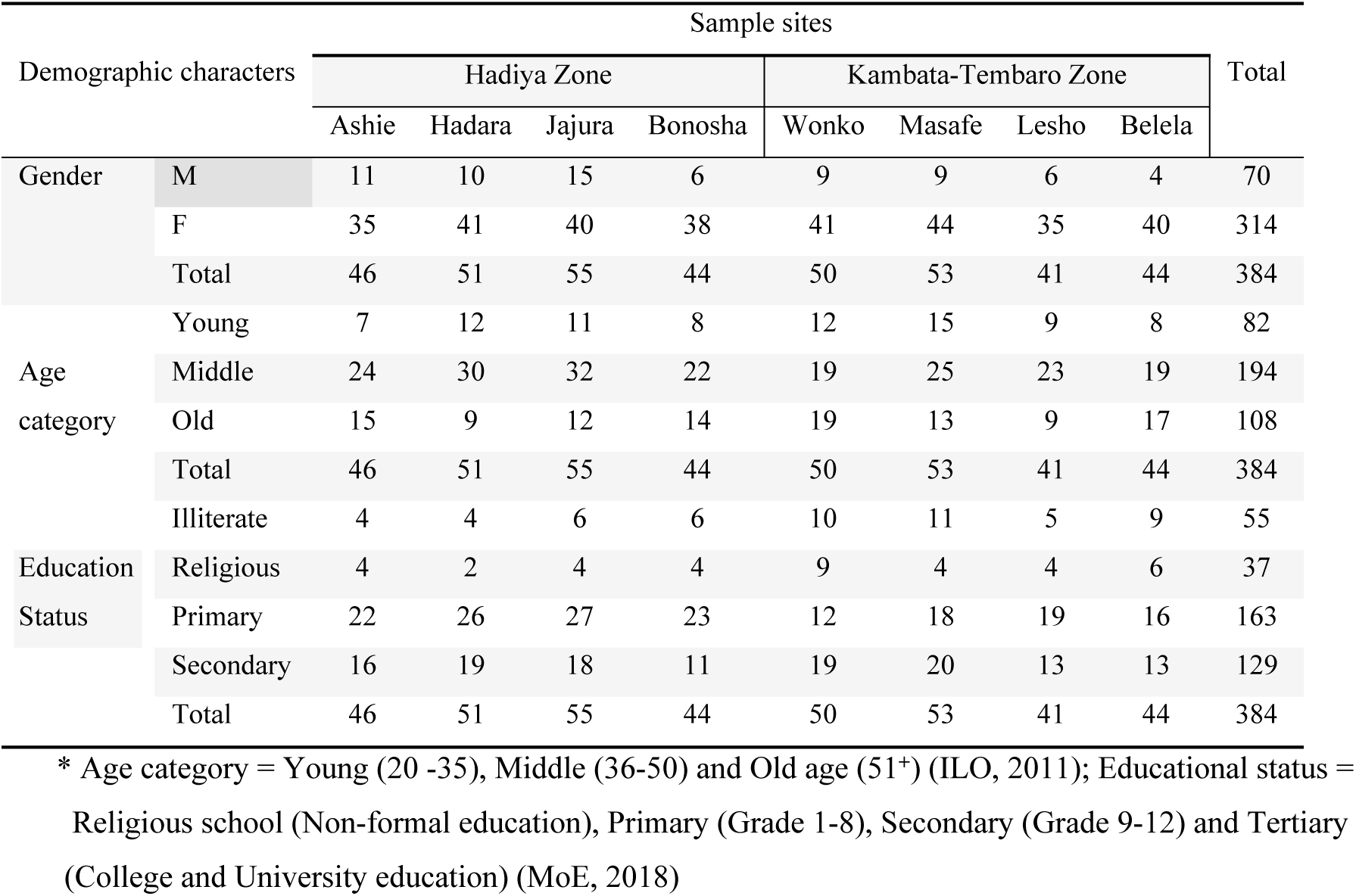
Respondents Profile

**Table 2:** provides a detailed listing of these spices, including their growth forms, vernacular (Amharic) names, and their occurrence in homegardens.

| S/No | Scientific Name | Family | Amharic Name | Habit |
| --- | --- | --- | --- | --- |
| 1 | *Aframomum corrorima* | Zingiberaceae | Kororima | Herbaceous |
| 2 | *Allium sativum* | Amaryllidaceae | Nech Shinkurt | Herbaceous |
| 3 | *Anethum foeniculum* | Apiaceae | Ensilal | Herbaceous |
| 4 | *Artemisia afra* | Asteraceae | Hintam | Shrub |
| 5 | *Artemisia rehan* | Asteraceae | Rehan | Herbaceous |
| 6 | *Brassica nigra* | Brassicaceae | Tikur Senafich | Herbaceous |
| 7 | *Brucea antidysenterica* | Simaroubaceae | Emismael | Shrub |
| 8 | *Camomum subulatum* | Zingiberaceae | K'amomum | Herbaceous |
| 9 | *Capsicum annuum* | Solanaceae | Berbere | Herbaceous |
| 10 | *Cinnamomum zeylanicum* | Lauraceae | Qerenfud/Kerenfud | Tree |
| 11 | *Citrus aurantifolia* | Rutaceae | Lomi Marany | Shrub/Tree |
| 12 | *Citrus limonia* | Rutaceae | Lomi Ambasah | Shrub/Tree |
| 13 | *Coriandrum sativum* | Apiaceae | Dimbilal | Herbaceous |
| 14 | *Curcuma longa* | Zingiberaceae | Ird | Herbaceous |
| 15 | *Cuminum cyminum* | Apiaceae | Kemen | Herbaceous |
| 16 | *Elsholtzia spp.* | Lamiaceae | Embamaarish | Herbaceous |
| 17 | *Foeniculum vulgare* | Apiaceae | Ensilal | Herbaceous |
| 18 | *Lippia abyssinica* | Verbenaceae | Koseret | Shrub |
| 19 | *Lippia adoensis* | Verbenaceae | Koseret | Shrub |
| 20 | *Nigella sativa* | Ranunculaceae | Tikur Azmud | Herbaceous |
| 21 | *Ocimum americanum* | Lamiaceae | Osim | Herbaceous |
| 22 | *Ocimum basilicum* | Lamiaceae | Besobila | Herbaceous |
| 23 | *Ocimum lamiifolium* | Lamiaceae | Demakese | Herbaceous |
| 24 | *Origanum vulgare* | Lamiaceae | Oregano | Herbaceous |
| 25 | *Osyris quadripartita* | Santalaceae | Emtaw | Shrub |
| 26 | *Piper nigrum* | Piperaceae | Kundo Berbere | Vine |
| 27 | *Rhamnus prinoides* | Rhamnaceae | Gesho | Shrub |
| 28 | *Rosmarinus officinalis* | Lamiaceae | Rozmari | Shrub |
| 29 | *Ruta chalepensis* | Rutaceae | Tenadam | Herbaceous |
| 30 | *Salvia officinalis* | Lamiaceae | Salviya | Herbaceous |
| 31 | *Syzygium aromaticum* | Myrtaceae | Kerenfuz | Flower bud/Tree |
| 32 | *Thymus vulgaris* | Lamiaceae | Tosina | Herbaceous |
| 33 | *Trachyspermum ammi* | Apiaceae | Ajowan | Herbaceous |
| 34 | *Trigonella foenum-graecum* | Fabaceae | Abish | Herbaceous |
| 35 | *Zingiber officinale* | Zingiberaceae | Zinjibil | Herbaceous |

**Table 3:** Spices Diversity Indices

| Occurrence | H' | D1 | D2 | RI |
| --- | --- | --- | --- | --- |
| Homegarden | 2.62 | 0.88 | 0.81 | 7.63 |
| Farmland | 1.68 | 0.69 | 0.71 | 11.92 |
| Wild area | 0.64 | 0.42 | 0.31 | 12.44 |

H' = Shannon diversity index, D1 = Simpson diversity index, D2 = Simpson evenness, RI = Richness index (Margalef index)

### 3.4. Ethnobotanical Roles of Spices

This study’s findings reveal the significant and interconnected roles that spices fulfill within the culinary traditions, medicinal systems, and ritual practices of the Hadiya and Kambata-Tembaro zones in Central Ethiopia (Table 4). These roles are indicative of both rich ecological knowledge and enduring cultural heritage, positioning spices as a crucial biocultural nexus that bridges environment, health, and identity. By situating these findings within the broader ethnobotanical literature, this discussion identifies both commonalities and unique regional expressions.

**Table 4:** Integrated Ethnobotanical Roles of Spices in the Hadiya and Kambata-Tembaro Area (Culinary uses, Medicinal values, and Ritual Applications

| Scientific Name | Culinary Use | Medicinal Values | Ritual Applications |
| --- | --- | --- | --- |
| *Aframomum corrorima* | Key in *berbere*, stews, spiced butter | Treats coughs, colds, digestion | Burned in incense, postpartum steam baths |
| *Allium sativum* | Universally used in sauces and meats | Antibacterial, cardiovascular health | Used in protective charms |
| *Anethum foeniculum* | Flavoring, herbal teas | Relieves bloating, aids lactation | Used in postpartum steam |
| *Artemisia afra* | Rarely culinary | Cold, malaria, digestive aid | Fumigation, cleansing spaces |
| *Artemisia rehan* | Spiced butter in some areas | Antiseptic, digestive aid | Used in smoke rituals and healing |
| *Brucea antidysenterica* | Not culinary | Treats dysentery, parasites | Associated with healing rituals |
| *Camomum subulatum* | Occasional tea flavoring | Sedative, pain relief | Used in purification baths |
| *Capsicum annum* | Vital in *berbere*, sauces | Stimulates appetite, circulation | Occasionally burned or ingested ritually |
| *Cinnamomum zeylanicum* | Used in drinks, sweets, meat dishes | Antimicrobial, anti-inflammatory | Incense, aromatic ceremonies |
| *Citrus aurantifolia* | Marinades, souring agent | Detoxification, throat soother | Cleansing rituals |
| *Citrus limonia* | Fresh consumption, beverages | Immunity and digestion booster | Minor ritual cleansing |
| *Coriandrum sativum* | Seeds in spice mixes | Bloating, mild laxative | Rarely ritual |
| *Cuminum cyminum* | Seasoning *shiro*, sauces | Digestion, gas relief | Rarely ritual |
| *Curcuma longa* | Coloring and flavoring stews | Anti-inflammatory, skin healing | Postpartum painting, ritual baths |
| *Elsholtzia spp.* | Herbal teas | Treats fever, headaches | Used in *tish* and fumigation |
| *Foeniculum vulgare* | Herbal teas | Gas relief, lactation stimulant | Occasionally used postpartum |
| *Hagenia abyssinica* | Not culinary | Powerful antiparasitic (tapeworm) | Consumed ceremonially once a year |
| *Lippia abyssinica* | Flavoring in butter, sauces | Cold remedy, appetite stimulant | Burned in postpartum rituals |
| *Lippia adoensis* | Mild spice for butter and drinks | Respiratory and stomach ailments | Steam and bath rituals |
| *Nigella sativa* | Baked goods, spice blends | Immune boosting, anti-inflammatory | Protective and blessing uses |
| *Ocimum americanum* | Stews and teas | Fever, colds, stomach issues | Smudging, spiritual baths |
| *Ocimum basilicum* | Spiced butter, sauces | Relieves stress, digestion | Used in steam healing rituals |
| *Ocimum lamiifolium* | Not culinary | Headache, fever, eye infection | Used in postpartum, child healing rituals |
| *Origanum vulgare* | Meat and sauce seasoning | Respiratory, antimicrobial | Used in fumigation |
| *Osyris quadripartita* | Rarely culinary | Bone fractures, gynecological aid | Used in ancestral rituals |
| *Piper nigrum* | Meat seasoning, sauces | Digestion, warming effect | Included in protective mixtures |
| *Rhamnus prinoides* | Fermentation of *t'ej*, *tella* | Mild laxative, digestion | Central in brewing rituals |
| *Rosmarinus officinalis* | Seasoning meat and stews | Circulation, memory stimulant | Incense in blessings and ceremonies |
| *Ruta chalepensis* | Spiced butter, drinks | Menstrual cramps, stomach pain | Wards off evil, ritual protection |
| *Salvia officinalis* | Mild butter or herbal use | Anti-inflammatory, sore throat | Used in cleansing smudges |
| *Syzygium aromaticum* | Drinks, sweets, spice blends | Toothaches, antiseptic | Burned for fragrance and blessing |
| *Thymus vulgaris* | Meat and soup flavoring | Cough relief, digestive aid | Used in ritual purification |
| *Trachyspermum ammi* | Rare spice mix use | Relieves gas, indigestion | Postpartum bath ingredient |
| *Trigonella foenum-graecum* | *Shiro*, *berbere*, porridge | Promotes lactation, blood sugar balance | Used in postpartum healing drinks |
| *Zingiber officinale* | Meats, drinks, spice blends | Nausea, inflammation relief | Healing rituals and tonics |

### 3.5. Culinary Preparation Techniques of Spices

Our findings (Table 5) provide comprehensive documentation of the culinary use of various spices in the study area, highlighting both the plant parts utilized and the preparation methods employed in local food traditions. The results reveal a rich body of indigenous culinary knowledge tied to plant diversity that a wide range of plant parts including seeds, fruits, leaves, bulbs, and rhizomes are utilized in local cuisine, depending on the species and the specific dish being prepared. Seeds and fruits are predominantly used for flavoring stews and sauces, while leaves and bulbs often serve as essential ingredients in the preparation of spice mixtures. Preparation techniques vary widely and include drying and grinding, roasting, crushing, boiling, and blending, each contributing to the unique flavor profiles of traditional dishes. Many of the spices are key components in staple culinary preparations such as *wot*, *berbere*, and *mitmita*, underscoring their integral role in the local food culture. Beyond flavor enhancement, these spices also fulfill functional roles as preservatives and digestive aids, reflecting the deep interconnection between culinary practice and traditional knowledge.

**Table 5:**
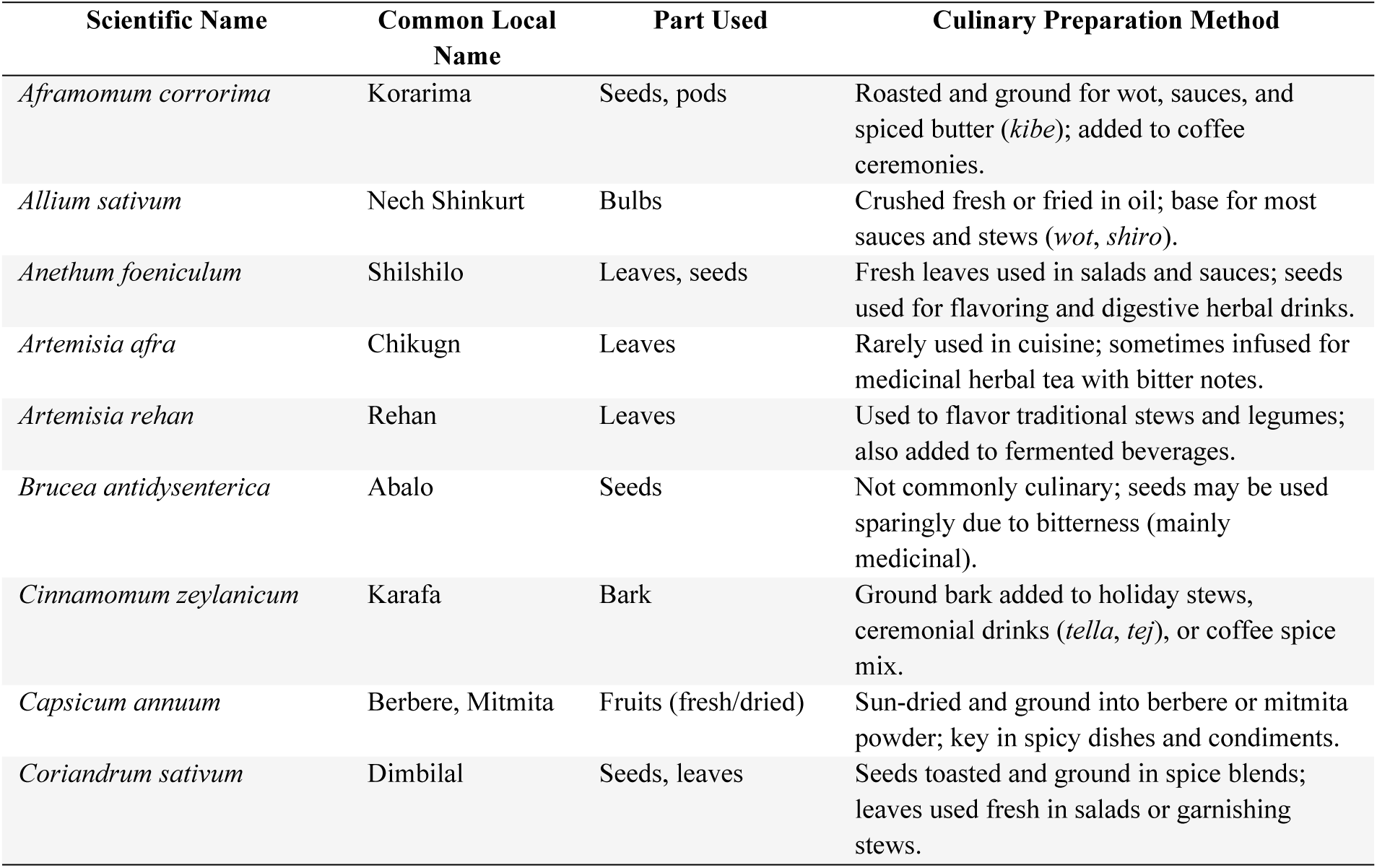
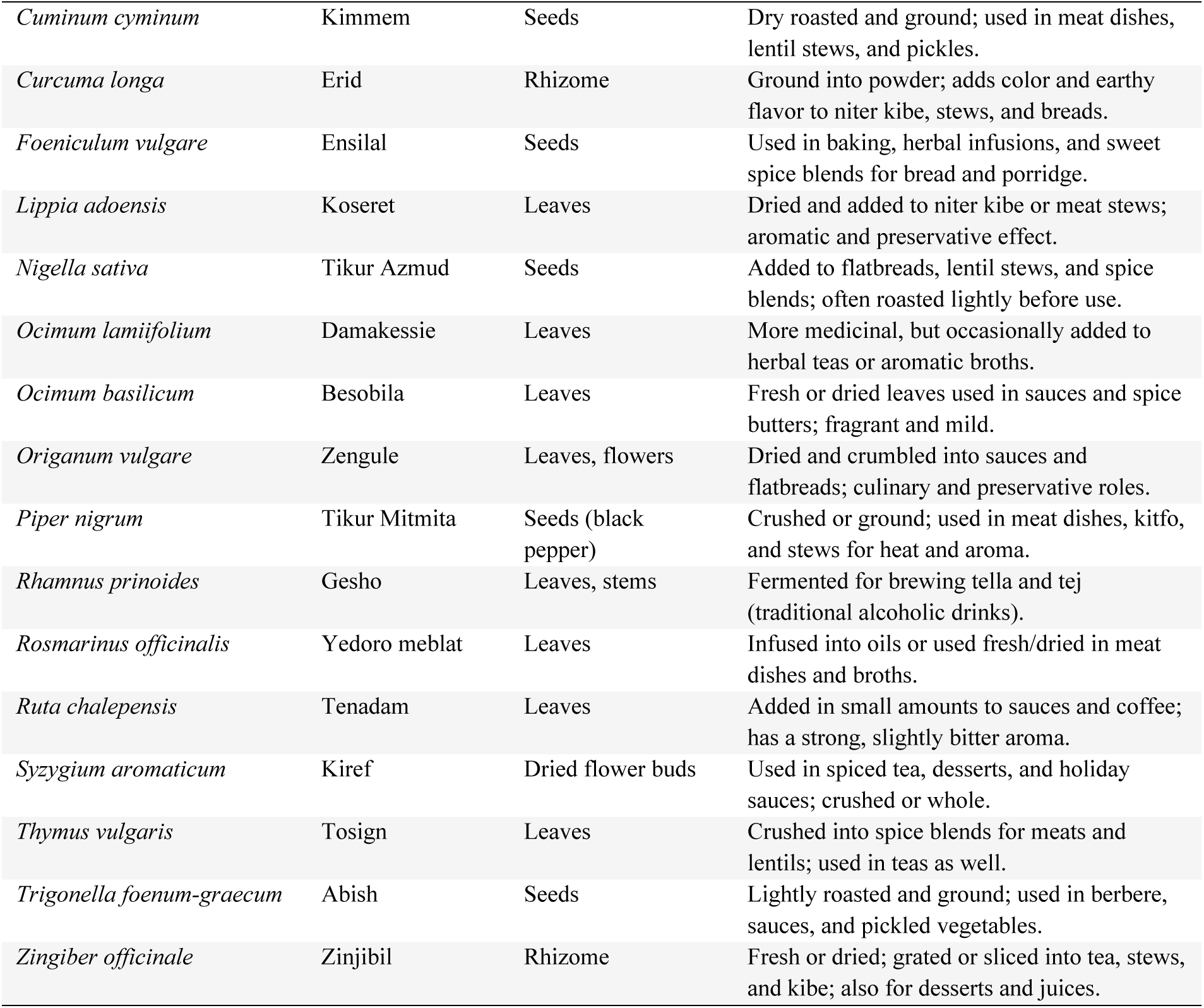
Overview of Spice Plant Parts utilized and Their Corresponding Culinary Preparation Techniques

| Scientific Name | Common Local Name | Part Used | Culinary Preparation Method |
| --- | --- | --- | --- |
| *Aframomum corrorima* | Korarima | Seeds, pods | Roasted and ground for wot, sauces, and spiced butter (*kibe*); added to coffee ceremonies. |
| *Allium sativum* | Nech Shinkurt | Bulbs | Crushed fresh or fried in oil; base for most sauces and stews (*wot*, *shiro*). |
| *Anethum foeniculum* | Shilshilo | Leaves, seeds | Fresh leaves used in salads and sauces; seeds used for flavoring and digestive herbal drinks. |
| *Artemisia afra* | Chikugn | Leaves | Rarely used in cuisine; sometimes infused for medicinal herbal tea with bitter notes. |
| *Artemisia rehan* | Rehan | Leaves | Used to flavor traditional stews and legumes; also added to fermented beverages. |
| *Brucea antidysenterica* | Abalo | Seeds | Not commonly culinary; seeds may be used sparingly due to bitterness (mainly medicinal). |
| *Cinnamomum zeylanicum* | Karafa | Bark | Ground bark added to holiday stews, ceremonial drinks (*tella*, *tej*), or coffee spice mix. |
| *Capsicum annuum* | Berbere, Mitmita | Fruits (fresh/dried) | Sun-dried and ground into berbere or mitmita powder; key in spicy dishes and condiments. |
| *Coriandrum sativum* | Dimbilal | Seeds, leaves | Seeds toasted and ground in spice blends; leaves used fresh in salads or garnishing stews. |
| *Cuminum cyminum* | Kimmem | Seeds | Dry roasted and ground; used in meat dishes, lentil stews, and pickles. |
| *Curcuma longa* | Erid | Rhizome | Ground into powder; adds color and earthy flavor to niter kibe, stews, and breads. |
| *Foeniculum vulgare* | Ensila | Seeds | Used in baking, herbal infusions, and sweet spice blends for bread and porridge. |
| *Lippia adoensis* | Koseret | Leaves | Dried and added to niter kibe or meat stews; aromatic and preservative effect. |
| *Nigella sativa* | Tikur Azmud | Seeds | Added to flatbreads, lentil stews, and spice blends; often roasted lightly before use. |
| *Ocimum lamifolium* | Damakessie | Leaves | More medicinal, but occasionally added to herbal teas or aromatic broths. |
| *Ocimum basilicum* | Besobila | Leaves | Fresh or dried leaves used in sauces and spice butters; fragrant and mild. |
| *Origanum vulgare* | Zengule | Leaves, flowers | Dried and crumbled into sauces and flatbreads; culinary and preservative roles. |
| *Piper nigrum* | Tikur Mitmita | Seeds (black pepper) | Crushed or ground; used in meat dishes, kitfo, and stews for heat and aroma. |
| *Rhamnus prinoides* | Gesho | Leaves, stems | Fermented for brewing tella and tej (traditional alcoholic drinks). |
| *Rosmarinus officinalis* | Yedoro meblat | Leaves | Infused into oils or used fresh/dried in meat dishes and broths. |
| *Ruta chalepensis* | Tenadam | Leaves | Added in small amounts to sauces and coffee; has a strong, slightly bitter aroma. |
| *Syzygium aromaticum* | Kiref | Dried flower buds | Used in spiced tea, desserts, and holiday sauces; crushed or whole. |
| *Thymus vulgaris* | Tosign | Leaves | Crushed into spice blends for meats and lentils; used in teas as well. |
| *Trigonella foenum-graecum* | Abish | Seeds | Lightly roasted and ground; used in berbere, sauces, and pickled vegetables. |
| *Zingiber officinale* | Zinjibil | Rhizome | Fresh or dried; grated or sliced into tea, stews, and kibe; also for desserts and juices. |

The study identified eight distinct traditional dishes that are commonly prepared using locally cultivated spices. These dishes including *Anaqaala*, *Ataakana*, and *Hanissabela* are primarily prepared by women, who serve as both caregivers and culinary experts within their households, locally referred to as *Ammo’o*. The consistent use of key spices such as *Allium sativum* (garlic), *Capsicum frutescens* (chili), and *Ruta chalepensis* (rue) not only enhances the flavor and aroma of these dishes but also contributes to their perceived medicinal value. These traditional preparations are deeply embedded in the local diet and serve as expressions of cultural identity, highlighting the important role of women in preserving and transmitting culinary knowledge across generations.

### 3.6. Community Preference Ranking of Spices

The results of the preference ranking exercise, as presented in Table 5, offer a clear depiction of the local communities’ prioritization of spices based on a multidimensional valuation system. Each spice was assigned a score reflecting its perceived importance, determined by factors such as frequency of use, culinary versatility, symbolic and ritual significance, and degree of cultural embeddedness in daily and ceremonial life. Higher scores indicate stronger local preference and wider cultural relevance, while lower scores correspond to more limited or specialized use.

Capsicum annuum (commonly known as *berbere* or *mitmita*) emerged as the most highly favored spice across both study zones. Its widespread use in everyday cooking, particularly as a base ingredient in iconic Ethiopian dishes such as doro wot, shiro, and kitfo, underscores its central role in culinary identity. Beyond flavoring, it also serves as a marker of hospitality and communal feasting, contributing to its symbolic value. The consistently high scores assigned to *Capsicum annuum* reflect both its culinary indispensability and its deep cultural resonance within local foodways. Other spices, such as *Aframomum corrorima* and *Allium sativum*, also ranked highly due to their dual use in flavoring and ritual or medicinal contexts, highlighting the multifunctionality of spice plants in Hadiya and Kambata-Tembaro traditions.

### 3.7. Conservation status and Threats to Spices

Fig. 2 presents the key threats identified by the community as impacting the sustainability and diversity of spices within the study area, categorized for clarity and based on the data collected through semi-structured interviews, focus group discussions, and preference ranking exercises. This figure synthesizes the findings from multiple data collection methods, offering a comprehensive understanding of the pressures these valuable plants face. The figure is organized around distinct Threat Categories, which emerged as recurring themes during the semi-structured interviews and focus group discussions. These categories provide a framework for understanding the multifaceted nature of the threats.

**Figure 2.**
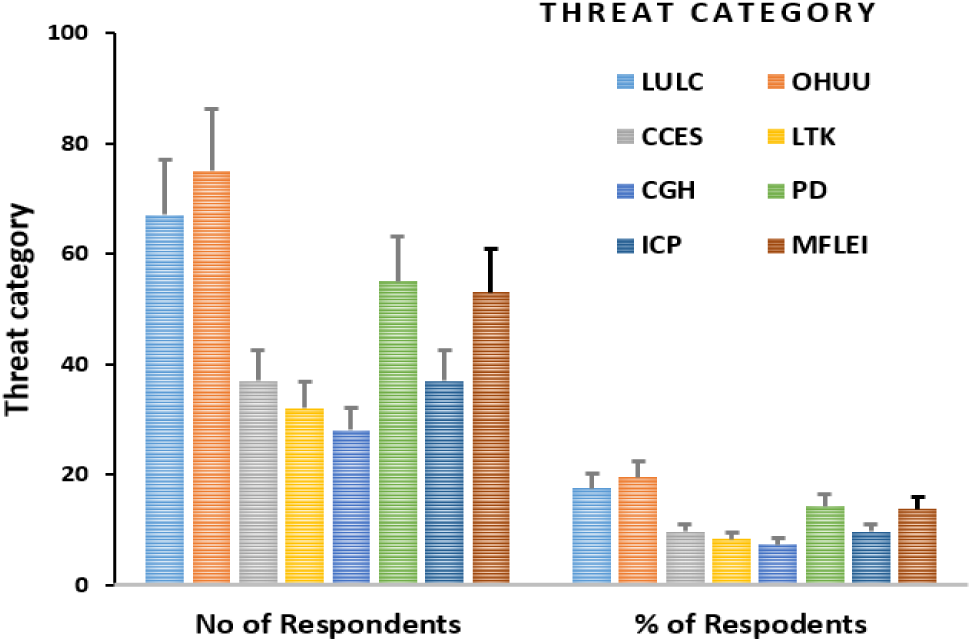
Threats category to Spices (LULC= Land-Use Change & Habitat Loss, OHUU= Overharvesting & Unsustainable Use, CCES= Climate Change & Environmental Stress, LTK= Loss of Traditional Knowledge, CGK= Commercialization & Genetic Homogenization, PD=Pests and Diseases, ICP=Inadequate Conservation Policies, MFLEI= Market Fluctuations & Low Economic Incentives).

### 3.7. Socioeconomic and Gendered Dimensions of Spice Livelihoods

Table 6 presents key findings on the gendered and socioeconomic roles in spice production and trade across the study area. The findings reveal that while both men and women are deeply involved in spice-related activities, there are significant inequalities in access to resources, participation in training, and control over income. For example, although 48% of women reported that spices are their main income source, only 12% owned the land used for spice cultivation. Access to capacity-building opportunities is similarly limited, with just 15% of women having received any form of training, compared to much higher participation by men. Additionally, in male-headed households, 77% of decisions regarding spice income were made by men, indicating a clear gender gap in financial control and autonomy.

**Table 6.** Results of preference ranking of spices based on frequency of use-values exercised by key informants (KI) and focus group discussants (FGD). Scores in the table indicate ranks given to multipurpose trees based on their use-values (highest number (10) was given for the tree species which informants thought most- preferred in its use-values, and the lowest number (1) was given for its least-preference). The highest total scores were the most-preferred tree species by the farmers. Letters A-J represent KI and FGD.

| Spices | Key informants and focus group discussants labeled A-J |  |  |  |  |  |  |  |  |  | Total score | Rank |
| --- | --- | --- | --- | --- | --- | --- | --- | --- | --- | --- | --- | --- |
|  | A | B | C | D | E | F | G | H | I | J |  |  |
| *Aframomum corrorima* | 8 | 7 | 6 | 10 | 9 | 7 | 10 | 9 | 6 | 8 | 80 | 2 |
| *Allium sativum* | 6 | 5 | 10 | 7 | 5 | 8 | 8 | 5 | 9 | 10 | 73 | 3 |
| *Coriandrum sativum* | 4 | 9 | 5 | 4 | 6 | 9 | 4 | 8 | 7 | 4 | 60 | 6 |
| *Ruta chalepensis* | 3 | 3 | 4 | 5 | 3 | 1 | 3 | 2 | 3 | 1 | 28 | 8 |
| *Nigella sativa* | 7 | 4 | 3 | 9 | 7 | 5 | 9 | 7 | 8 | 7 | 63 | 5 |
| *Lippia adoensis* | 5 | 6 | 9 | 6 | 4 | 4 | 5 | 4 | 4 | 5 | 52 | 7 |
| *Zingiber officinale* | 9 | 8 | 7 | 3 | 8 | 6 | 6 | 6 | 5 | 6 | 64 | 4 |
| *Capsicum annuum* | 10 | 10 | 8 | 8 | 10 | 10 | 7 | 10 | 10 | 9 | 92 | 1 |
| *Curcuma longa* | 2 | 1 | 1 | 2 | 2 | 3 | 2 | 3 | 2 | 2 | 20 | 9 |
| *Ocimum basilicum* | 1 | 2 | 2 | 1 | 1 | 2 | 1 | 1 | 1 | 3 | 15 | 10 |

**Table 6:** Socioeconomic and Gendered Dimensions of Spice

| Parameter | Findings | Gender Disaggregation |
| --- | --- | --- |
| Household Income from Spices | 62% of households earn >40% of annual income from spice-related activities. | 48% of women say spice is their main income source. |
| Market Participation | Majority sell at local markets; limited access to regional buyers or value-added chains. | Men more involved in transport and sales. |
| Access to Land | 76% of respondents cultivate spices on small plots (<0.5 ha). | Only 12% of women own land used for spice farming. |
| Access to Training | 35% of respondents have attended spice-related trainings (agronomy, processing). | Only 15% of trainees were women. |
| Division of Labor | Women: planting, weeding, drying, storage.<br>Men: land prep, marketing. | Labor is heavily gendered. |
| Control Over Income | In male-headed households, 77% of income decisions made by men. | Only 23% of women control spice income they earn. |
| Involvement in Cooperatives | 40% of respondents are members of spice-related cooperatives. | Women make up only 18% of cooperative members. |
| Decision-Making Power | Men are primary decision-makers on land use, marketing, and investments. | Women report limited influence on economic decisions. |
| Time Burden & Constraints | Women cite lack of time due to domestic responsibilities as a major barrier to training or market engagement. | High time poverty among women. |
| Emerging Opportunities | Women-led microenterprises in fenugreek and turmeric processing noted in Hosaena and Durame. | Early signs of economic empowerment for women. |

The data also show that women are underrepresented in formal structures such as cooperatives, with only 18% membership, despite their critical role in cultivation and post-harvest processing. These disparities reflect broader social norms and structural barriers that limit women’s economic empowerment. However, the study also identified promising developments, particularly in Hadiya and Kembata-Tembaro areas, where women have begun forming small-scale enterprises in fenugreek and turmeric processing. These emerging trends suggest potential pathways for enhancing women’s roles in the value chain. Overall, the results underscore the importance of designing inclusive programs that ensure equitable access to land, training, markets, and decision-making processes for marginalized groups, especially women.

## 4. Discussion

### 4.1. Demographic Profiles of the Respondents

The demographic composition of the respondents provides important context for interpreting the ethnobotanical knowledge documented in the study. A total of 384 individuals participated, comprising 314 females and 70 males. The significantly higher representation of female respondents reflects the central role women play in spice cultivation, usage, and preservation in the study area. This gendered knowledge distribution is consistent with previous ethnobotanical research, which has shown that women are often the primary custodians of plant-based knowledge, particularly in homegarden systems (Howard, 2003; Ghimire et al., 2004).

Respondents ranged in age from 20 to 67 years, with a mean age of 43.45 years (±9.95 SD), indicating that the knowledge shared spans multiple generations. This age diversity enhances the richness of insight into both inherited traditions and contemporary practices regarding spice use. Educationally, 42.4% of participants had completed primary school and 33.6% had attained secondary education. Moderate levels of formal education among respondents may influence the way traditional knowledge is preserved and transmitted, as well as how it is integrated with modern practices (Zent, 2009).

The average household size was 5.76 individuals (±2.24 SD), with 89.1% of households comprising 4 to 7 members. The largest family size recorded was 9, observed in all study sites except Ashie. Larger family units may support a more diverse range of agricultural activities, including the cultivation of various spice species, while also facilitating intergenerational knowledge transfer an essential component of ethnobotanical knowledge systems (Reyes-García et al., 2005).

### 4.2. Spices in the study area

The documentation of 35 spice and condiment species belonging to 27 genera and 17 families in Hadiya and Kambata- Tembaro underscores the region’s rich ethnobotanical heritage and its role as a center of traditional knowledge and plant-based utility. The predominance of herbaceous species (63%) reflects a broader trend in Ethiopian ethnobotany, where such species are preferred due to their short growth cycles, ease of propagation, and adaptability to household agroecosystems (Addis et al., 2005; Giday et al., 2009). Herbaceous plants also tend to be more accessible to women and elders, who are often the primary custodians of traditional culinary and medicinal knowledge (Kassaye et al., 2006).

The taxonomic distribution, led by the family *Lamiaceae* (8 species), aligns with global patterns where this family is recognized for its aromatic members, rich in essential oils and bioactive compounds (Van Wyk & Wink, 2004).

Species within *Lamiaceae*, such as *Ocimum* and *Thymus*, play a central role in Ethiopian spice mixtures and traditional medicine, which reinforces their local importance. Similar findings were reported by Jansen (1981) and Lulekal et al. (2011), who noted the widespread culinary and therapeutic relevance of Lamiaceae species in other Ethiopian regions.

The high incidence of spices grown in homegardens (80%) confirms that these micro-agroecosystems are key in maintaining both plant diversity and ethnobotanical knowledge (Berkes et al., 2000; Hamilton, 2004). Homegardens are traditionally managed by women and elders, who ensure the transmission of knowledge across generations through informal channels (Posey, 1999; Pankhurst, 2001). This aligns with findings from southwestern Ethiopia and other highland regions where homegardens support ethnomedicinal species and play a role in in-situ conservation (Giday et al., 2007).

The diverse uses of spices (culinary, medicinal, and ritual) highlight the multifunctionality of these plants in local livelihoods. For instance, spices such as *Aframomum corrorima*, *Artemisia afra*, and *Nigella sativa* are not only dietary additives but also integral to healing practices and ceremonial life. This resonates with the concept of “bio-cultural keystone species” (Garibaldi & Turner, 2004), which are culturally significant plants central to identity, tradition, and well-being.

Additionally, the detailed vernacular taxonomy and morphological identification practices observed in the study area point to an advanced and nuanced local classification system, often diverging from Western Linnaean taxonomy but deeply rooted in practical knowledge and sensory cues (Berlin, 1992). Such indigenous classification systems contribute significantly to ethnobiological resilience and the conservation of traditional ecological knowledge (TEK).

Comparatively, the results echo earlier ethnobotanical studies from other parts of Ethiopia, such as those among the Meinit (Giday et al., 2007) and Zay communities (Giday et al., 2009), which similarly emphasize the intricate relationships between biodiversity, culture, and livelihood. However, this study brings novel regional insights by focusing on the central highlands of Hadiya and Kambata-Tembaro, areas previously underrepresented in ethnobotanical research. The observed pattern of spice use also provides a counter-narrative to the often commodity-driven perception of spices, instead illustrating their embeddedness in everyday health, ritual, and culinary practices.

Ultimately, these findings reinforce the need for inclusive conservation policies that respect and integrate local knowledge systems. Recognizing the interwoven cultural and ecological dimensions of spices could enhance both biodiversity conservation and food security initiatives in Ethiopia and beyond.

### 4.3. Diversity of spices

Our findings reveal a clear gradient in spice species diversity across the investigated land-use systems, with homegardens exhibiting significantly higher Shannon and Simpson diversity indices compared to farmland and the wild area. The high spice diversity observed in homegardens aligns with existing literature highlighting the role of these agroforestry systems as biodiversity reservoirs, particularly for useful plant species (Kumar & Nair, 2004; Fernandes & Nair, 1986). Homegardens, characterized by their multi-strata structure and intensive management practices, often integrate a wide array of plant species to meet diverse household needs, including culinary, medicinal, and ornamental purposes (Soemarwoto, 1987). The deliberate cultivation and integration of various spice and condiment species within homegardens contribute to the observed high species richness and evenness. This intentional diversification acts as a form of *in situ* conservation, safeguarding a broader range of genetic resources compared to more monoculture-oriented systems like farmland.

The lower spice diversity in farmland (Shannon: 1.68, Simpson: 0.69) is likely a consequence of agricultural intensification and a focus on a limited number of primary crops. Modern agricultural practices often prioritize high-yielding varieties of staple crops, leading to the simplification of agroecosystems and the displacement of associated biodiversity, including spices and condiments (Perfecto et al., 1992). While some spice species might be cultivated as intercrops or in boundary plantings, their diversity and evenness are generally lower compared to the integrated and multi-functional nature of homegardens. Our findings corroborate studies that have documented reduced biodiversity in agricultural landscapes compared to more traditional and diversified farming systems (Altieri, 1999).

The significantly lower spice diversity in the wild area (Shannon: 0.64, Simpson: 0.42) presents an interesting contrast. While one might intuitively expect wild habitats to harbor high biodiversity, our results suggest a less even distribution and lower richness of spice species in the specific wild area investigated. This could be attributed to several factors. Firstly, the “wild area” as defined in our study might represent a specific ecological niche or habitat type that naturally supports a limited range of spice species. Secondly, factors such as habitat degradation due to external pressures ( encroachment, resource extraction) or natural environmental limitations could contribute to lower diversity and evenness. It is also possible that the sampling effort in the wild area captured a specific patch with lower spice concentration. These findings underscore the importance of clearly defining and characterizing “wild areas” in biodiversity studies, as they are not always inherently more diverse for all plant groups compared to managed systems. Similar findings of lower diversity for specific plant groups in certain wild habitats compared to managed agroecosystems have been reported in other studies (Bharucha & Pretty, 2010, on tree diversity).

The higher evenness of spice species in homegardens, as indicated by the relatively smaller difference between Shannon and Simpson indices (Table 3), suggests a more equitable distribution of abundance among the different spice species present. This contrasts with farmland and the wild area, where a few dominant species might account for a larger proportion of the observed spice occurrences, leading to lower evenness. This even distribution in homegardens could be a result of conscious selection and management by households to ensure a consistent supply of a variety of spices for different uses.

In conclusion, our findings strongly support the role of homegardens as crucial reservoirs of spice species diversity within the study area. The higher richness and evenness observed in these systems highlight their importance for *in situ* conservation and the maintenance of agrobiodiversity. The lower diversity in farmland underscores the potential negative impacts of agricultural intensification on associated plant diversity, while the findings from the wild area necessitate further investigation into the specific ecological and anthropogenic factors shaping its spice species composition. These results have significant implications for conservation strategies aimed at preserving spice diversity, suggesting a need to focus on supporting and promoting homegarden systems alongside sustainable management practices in both farmland and wild habitats.

### 4.4. Integrated Ethnobotanical Roles of Spices

The findings from this study illuminate the intricate and interdependent roles that spices play in the culinary traditions, medicinal systems, and ritual practices of the Hadiya and Kambata-Tembaro zones in Central Ethiopia. These roles not only reflect deep ecological knowledge and cultural heritage but also position spices as a biocultural nexus linking environment, health, and identity. This discussion situates the findings within broader ethnobotanical literature and identifies both convergences and unique regional dimensions.

#### Culinary Versatility and Regional Identity

The widespread culinary use of spices such as *Aframomum corrorima*, *Capsicum annuum*, *Allium sativum*, and *Zingiber officinale* in the study area aligns with patterns reported in other Ethiopian regions, where spices form the backbone of national dishes like *wot*, *doro*, and *shiro* (Jansen, 1981; Giday et al., 2009). However, the study area demonstrates a distinctive emphasis on the use of endemic or semi-endemic species, *Lippia abyssinica*, *Ocimum basilicum*, and *Ruta chalepensis* in the flavoring of local *niter kibbeh* (spiced clarified butter), tea infusions, and fermented beverages. This suggests a localized adaptation of pan-Ethiopian spice culture, shaped by ecological availability and cultural preference.

The inclusion of lesser-known taxa such as *Elsholtzia spp.* and *Camomum subulatum* in culinary applications also reflects a potential underreporting of spice species in past literature (Abebe & Ayehu, 1993), and contributes to the call for finer-scale documentation of local food ethnobotanies.

#### Medicinal Functions and Ethnopharmacological Continuity

The medicinal roles of spices in the study area resonate with extensive ethnopharmacological findings across Ethiopia and the Horn of Africa. For instance, the use of *Nigella sativa*, *Curcuma longa*, and *Trigonella foenum-graecum* in managing inflammation, digestive disorders, and postnatal care is well documented (Giday et al., 2007; Teklehaymanot & Giday, 2007). However, this study expands on this knowledge by identifying nuanced medicinal uses that are deeply embedded in local life stages and social structures such as postpartum healing (*dukana*) practices involving *Ocimum lamiifolium*, *Anethum foeniculum*, and *Lippia adoensis*.

The inclusion of ritualized medicine where medicinal herbs double as agents of spiritual cleansing or protection is consistent with the concept of “medical pluralism” observed in other African ethnomedical systems (Van Wyk & Wink, 2004). Particularly, the dual use of *Artemisia afra*, *Ruta chalepensis*, and *Osyris quadripartita* as both pharmacological and spiritual agents challenges the biomedical compartmentalization of plant uses and calls for an integrated understanding of local health epistemologies.

#### Ritual Significance and Cultural Symbolism

Spices such as *Aframomum corrorima*, *Lippia abyssinica*, and *Ocimum species* are heavily embedded in ritual contexts in the study area from childbirth purification ceremonies and fumigation to healing incantations and protective smudging. This ritual use echoes studies from southern and southwestern Ethiopia, where spices and aromatic herbs are central to transitional rituals and ancestral veneration (Kassaye et al., 2006; Pankhurst, 2001).

What is notable in Hadiya and Kambata-Tembaro is the *fluid boundary* between ritual and therapeutic functions. For example, postpartum steaming using a complex mixture of spices serves medicinal, hygienic, and spiritual purposes simultaneously a finding that supports the holistic model of health in indigenous cosmologies (Berhane & Fahmi, 2010). The multifunctionality of spices here underlines their value not only as material resources but also as carriers of intangible heritage.

#### Comparative Ethnobotany and Cultural Specificity

While several of the spices identified in this study are widely used across Ethiopia, the distinct combinations, modes of application, and meanings attributed to them highlight the cultural specificity of spice ethnobotany. For instance, *Rhamnus prinoides* (used in fermenting *t’ej* and *tella*) is present in many regions, but the ceremonial protocols of its use and symbolic interpretations differ across communities. In the study area, its role extends beyond brewing into communal identity marking, indicating the need to investigate not just species presence, but their embeddedness in local worldviews.

Moreover, this study contributes new data on species such as *Brucea antidysenterica* and *Elsholtzia spp.*, which are underrepresented in national ethnobotanical inventories, despite having both medicinal and ritual significance. These findings validate the hypothesis that Ethiopia’s spice diversity is not only botanical but deeply cultural (Addis et al., 2005).

#### Implications for Biocultural Conservation

The observed convergence of culinary, medicinal, and ritual roles within single species points to the risk of “biocultural erosion” should these plants or their knowledge systems become marginalized. Spices in the study area are often conserved through active use, managed in home gardens, and transferred via oral tradition a pattern reported elsewhere in East Africa (Hamilton, 2004; Lulekal et al., 2011). However, threats such as land-use change, youth disinterest, and market commodification pose serious challenges. Integrating indigenous knowledge into local development plans and conservation policies, as advocated by scholars such as Posey (1999) and Berkes et al. (2000), is not only a cultural imperative but also an ecological one. Spices, as shown in this study, represent a convergence point where sustainable food systems, traditional medicine, and cultural resilience meet.

Generally, this study underscores the centrality of spices in the cultural and ecological lifeworlds of the Hadiya and Kambata-Tembaro people. By mapping their culinary, medicinal, and ritual applications, the research contributes to the growing field of ethnobotany that values the integrated, dynamic, and contextual nature of plant use. The findings affirm that spices are not merely flavoring agents or folk remedies, but critical vectors of knowledge, identity, and survival.

### 4.5. Culinary Preparation Techniques of Spices

The culinary application of spices in the study area reflects a profound integration of biodiversity, traditional knowledge, and cultural identity. Our findings (Table 5) document the extensive use of various plant parts including seeds, fruits, leaves, bulbs, and rhizomes in local food traditions. This diversity of plant parts corresponds to the multifunctionality of spices in culinary systems, serving not only to enhance flavor and aroma but also to improve texture, extend shelf life, and provide therapeutic benefits (Kuhnlein & Receveur, 1996; Pieroni & Price, 2006).

Seeds and fruits were found to be the most frequently used parts in the preparation of stews and sauces, particularly in the form of dried and ground powders. Leaves and bulbs, by contrast, were often used fresh or crushed to form aromatic bases for spice blends such as *berbere* and *mitmita*, which are essential in Ethiopian cuisine. These preparation methods including drying, roasting, boiling, and crushing demonstrate the community’s nuanced understanding of how different techniques influence flavor, potency, and storage life. Similar processing practices have been documented in other ethnobotanical studies, where traditional culinary methods serve to preserve not only the physical integrity of spices but also their bioactive properties (Oryan et al., 2021).

The study also identified eight culturally significant traditional dishes, including *Anaqaala*, *Ataakana*, and *Hanissabela*, which are deeply rooted in local dietary habits and frequently prepared using homegrown spices. These dishes are primarily prepared by women, known locally as *Ammo’o*, who hold specialized knowledge in food preparation, spice blending, and ritual practices associated with cooking. This finding aligns with prior ethnographic literature emphasizing the gendered dimensions of culinary knowledge and the role of women as primary custodians of traditional plant-based knowledge (Howard, 2003; Shiva, 1997).

The spices most frequently mentioned *Allium sativum* (garlic), *Capsicum frutescens* (chili), and *Ruta chalepensis* (rue) play a dual role in flavoring and health. Their widespread use is not merely for taste but also for their functional properties as digestive aids, antimicrobial agents, and traditional remedies for colds and stomach ailments. This dual use reflects the broader ethnobotanical principle of food-medicine continuum, where culinary plants serve both nutritional and therapeutic functions (Etkin & Ross, 1982; Pieroni & Quave, 2006).

The persistence of such practices in the face of changing dietary patterns and market integration suggests the resilience and adaptability of local knowledge systems. Moreover, the centrality of spice use in everyday meals and ceremonial contexts emphasizes the symbolic and social roles these plants play, reinforcing group identity and intergenerational continuity (Johns & Sthapit, 2004). As such, the culinary application of spices in the study area is not merely a matter of taste but a culturally embedded practice that encapsulates ecological knowledge, social structure, and heritage.

### 4.6. Community preference ranking of spices

The results of the preference ranking exercise provide critical insights into the socio-cultural dynamics and ethnobotanical valuation of spice species in Hadiya and Kambata-Tembaro zones. The multidimensional criteria used by local participants frequency of use, culinary versatility, symbolic resonance, and embeddedness in both daily life and ceremonial contexts mirror the broader conceptual frameworks in ethnobotanical literature that emphasize the cultural salience and functional diversity of plant species (Phillips & Gentry, 1993; Albuquerque et al., 2006).

*Capsicum annuum*, ranked as the most preferred spice, exemplifies this cultural salience. As observed in other Ethiopian ethnobotanical studies (Giday et al., 2009; Yineger & Yewhalaw, 2007), *Capsicum annuum* occupies a central position in both the flavor architecture of traditional dishes and in social performance rituals such as communal feasts, weddings, and religious observances. Its use in major national dishes like *doro wot* and *shiro* as well as its association with spice mixtures such as *berbere* and *mitmita* reinforces its status as a cornerstone of the Ethiopian culinary identity (Jansen, 2005; Mesfin, 2007). Additionally, its high demand and continuous cultivation in homegardens point to its economic and agronomic relevance.

Other highly ranked species such as *Aframomum corrorima* and *Allium sativum* illustrate the intersecting culinary, medicinal, and ritual values that define spice usage in the region. *Aframomum corrorima*, an indigenous spice also documented in the Gamo and Kaffa zones (Bahru et al., 2014), is not only a flavoring agent in ceremonial dishes and spiced butter (*niter kibe*) but is also employed in coffee rituals and healing practices, underlining its symbolic and therapeutic role. Similarly, *Allium sativum* is widely used for its antimicrobial properties (Teklehaymanot & Giday, 2007), while also serving as a staple in sauces and traditional broths, showcasing its functional duality.

The high preference scores attributed to such multifunctional spices align with findings from other ethnobotanical studies across Ethiopia and East Africa, which report that species with broader utility across culinary, medicinal, and ritual contexts tend to be more culturally valued and better conserved (De Boer & Lamxay, 2009; Asfaw, 2001). This supports the argument that preference ranking is not merely a reflection of taste or familiarity but also an indicator of integrated ecological knowledge and cultural identity.

Interestingly, some spices with limited use in ritual or medicinal settings, such as *Cinnamomum zeylanicum* or *Piper nigrum*, received moderate scores, possibly due to their imported status or restricted availability in local agroecosystems. This aligns with research that suggests locally accessible and culturally entrenched species are more highly prioritized than globally traded but culturally peripheral spices (Kujawska et al., 2015).

In sum, the preference rankings not only reveal the practical and symbolic significance of spices in the daily lives of the Hadiya and Kambata-Tembaro communities but also echo broader patterns observed in traditional societies where plant multifunctionality, cultural familiarity, and ecological access intersect to shape use priorities. These findings emphasize the importance of preserving local ethnobotanical knowledge, particularly in the face of cultural homogenization and agricultural modernization that may marginalize traditional preferences

### 4.7. Socioeconomic and Gendered Dimensions

The findings of this study provide a nuanced understanding of the socioeconomic and gendered roles embedded within the spice value chains in southern Ethiopia. The data show that while spices contribute significantly to rural household incomes (especially among smallholders) women, despite being heavily involved in labor-intensive aspects such as cultivation, harvesting, drying, and storage, continue to face structural barriers in accessing land, training, cooperative networks, and decision-making power. These observations align with earlier studies that have documented gender disparities in agricultural systems, where women contribute substantially to production but remain marginalized in terms of benefits and agency (Quisumbing et al., 2014; Farnworth et al., 2013).

The gendered division of labor identified in this study confirms patterns observed by Doss (2001), who noted that women tend to be concentrated in low-return and labor-intensive tasks within agricultural value chains, with men dominating more profitable roles such as marketing and transportation. The limited land ownership by women (12% in this study) is consistent with national-level analyses which show that women in Ethiopia rarely hold title deeds, often due to patriarchal inheritance practices and legal awareness gaps (Bezu & Holden, 2014). The lack of access to land directly impedes women’s eligibility for loans, extension services, and market-based contracts, reinforcing a cycle of exclusion.

Moreover, access to agricultural training remains highly unequal. Only 15% of women in the present study had received training related to spice agronomy or processing, echoing findings by World Bank (2015), which show that women are disproportionately underrepresented in formal capacity-building programs in sub-Saharan Africa. This not only limits women’s capacity to innovate and improve productivity but also affects the quality of spices entering the value chain. As such, low access to training and technology remains a critical constraint on enhancing both gender equity and product competitiveness.

Control over income is another significant area of disparity. The study revealed that in 77% of households, men were the sole decision-makers regarding the use of spice-related earnings. This is consistent with findings by Meinzen-Dick et al. (2011), who emphasized that even when women contribute labor or generate income, cultural norms often restrict their control over financial resources. This has direct implications for household well-being and nutrition, as women’s control over income is often associated with increased spending on food, health, and children’s education (Smith et al., 2003).

However, the emergence of women-led microenterprises in fenugreek and turmeric processing is a promising development, showing potential for transformative change. These grassroots initiatives reflect the broader trend of rural women engaging in value-added production as a means of circumventing traditional gender barriers. Similar trends have been documented in Uganda and India, where women’s engagement in processing and niche marketing of spices has increased their income and bargaining power (Njuki et al., 2011; Bhattarai et al., 2020).

This study also underscores the role of social capital and informal networks. Women, although underrepresented in formal cooperatives, often rely on informal trade relationships and women’s groups, which can offer alternative pathways for support and knowledge exchange. Building on this foundation through targeted support, such as women-led cooperative models or tailored training programs, could promote inclusive and sustainable growth of the spice sector.

In summary, the study confirms and extends the existing literature on gendered value chains, showing that systemic barriers continue to limit women’s full participation and benefit-sharing in spice economies. Addressing these gaps requires integrated strategies that combine asset redistribution (land rights), service accessibility (training and credit), and empowerment interventions (leadership and decision-making spaces). Doing so is not only a matter of gender justice but also of enhancing productivity, value chain efficiency, and community resilience.

Finally, the findings of this study have several important policy and ethical implications, particularly in relation to the integration of indigenous ecological knowledge (IEK) into national biodiversity conservation and food security agendas. The socioeconomic and gendered dimensions of spice livelihoods reveal that traditional knowledge, especially as held and transmitted by women plays a crucial role in sustaining agrobiodiversity, ensuring food security, and supporting rural economies. These findings reinforce arguments in the literature that indigenous knowledge systems are not only reservoirs of ecological wisdom but also strategic assets for climate resilience and sustainable agriculture (Berkes et al., 2000; Altieri & Toledo, 2011).

However, the undervaluation of this knowledge in formal policy frameworks remains a key challenge. Despite the significant contribution of women to spice cultivation, seed selection, and preservation practices, their knowledge is often excluded from institutional agricultural research and development processes. This exclusion not only reinforces existing gender inequalities but also undermines the potential of indigenous systems to inform sustainable development. As argued by Agarwal (1997), gendered ecological knowledge must be systematically recognized and incorporated into policy if conservation efforts are to be both effective and just.

Furthermore, the study raises critical questions around intellectual property rights (IPRs) and the ethics of knowledge extraction. As the commercial value of spices and traditional formulations increases, so does the risk of biopiracy— the unauthorized appropriation of community-held biological and cultural resources for commercial gain. This phenomenon has been widely documented in other regions, such as the neem and turmeric patent disputes in India (Shiva, 2007). Without appropriate legal safeguards, there is a real risk that indigenous communities—particularly women, who are often the custodians of such knowledge—may be excluded from the benefits derived from their own practices and innovations.

Accordingly, this study supports growing calls for community ownership and protection of biocultural heritage, as outlined by Posey & Dutfield (1996). National policies must incorporate provisions for free, prior, and informed consent (FPIC), community-defined benefit-sharing mechanisms, and the recognition of customary rights. For instance, the implementation of Access and Benefit Sharing (ABS) frameworks under the Nagoya Protocol (CBD, 2011) should be localized and operationalized with gender-sensitive tools to ensure that women’s contributions to spice-based knowledge systems are properly acknowledged and rewarded.

Moreover, ethical engagement with indigenous communities requires moving beyond extractive research paradigms toward participatory and reciprocal models of knowledge co-production. Researchers and policymakers must ensure that community members are not only consulted but are active collaborators and beneficiaries of biodiversity and food system interventions. This ethical commitment is particularly essential when working with marginalized or underrepresented groups, whose knowledge may otherwise be mined without compensation or recognition.

In conclusion, this study underscores the urgency of creating inclusive, equitable, and ethically sound policies that recognize indigenous knowledge as a living and evolving system. Safeguarding this knowledge through legal protection, community empowerment, and gender-responsive institutional frameworks is vital for both cultural survival and ecological sustainability. By elevating the voices of rural women and local knowledge holders, future biodiversity and food security strategies can become more socially just and ecologically effective.

## 5. Conclusion

This study presents a thorough ethnobotanical exploration of spice diversity in the Hadiya and Kambata-Tembaro zones of Central Ethiopia, uncovering a vibrant repository of traditional knowledge embedded in local cultures and livelihoods. The demographic composition of respondents marked by a significant involvement of women and representation across age groups reinforces the pivotal role women play as knowledge bearers and transmitters within their communities. The identification of 35 spice species, predominantly herbaceous and with the Lamiaceae family featuring prominently, underscores the area’s rich plant diversity. Homegardens emerged as vital ecological niches, serving not only as sources of sustenance but also as hubs of in situ conservation and intergenerational knowledge transfer. The multifunctional uses of spices in culinary practices, traditional medicine, and cultural rituals position these plants as biocultural keystone species, essential to the region’s social fabric. The observed variation in spice species across homegardens, farmlands, and wild areas further highlights the ecological importance of diverse land-use systems in sustaining agrobiodiversity and cultural heritage.

Equally important, the study sheds light on community preferences, culinary preparation methods, and the cultural logic underpinning spice selection, revealing complex valuation systems that blend sensory attributes with health beliefs and symbolic meanings. It also draws attention to the gendered dynamics and socioeconomic disparities within spice value chains, where women’s indispensable contributions are often underrecognized and underrewarded. These findings point to the necessity for inclusive, gender-sensitive, and ethically grounded policy interventions that respect and promote indigenous ecological knowledge particularly that held by women. Legal protections, community empowerment, and the equitable sharing of benefits must be central to biodiversity conservation and food security strategies. Looking forward, it is imperative to support longitudinal and participatory research focused on the ecological sustainability of spice species under shifting environmental conditions. Collaborative efforts between scientists, policymakers, and local communities especially women and elders are essential to preserving this ethnobotanical heritage. Such integrated approaches can ensure not only the conservation of spice biodiversity but also the empowerment of the communities who have nurtured and safeguarded this knowledge for generations.

